# Dominant species determines *Drosophila*-parasitoid food web dynamics and stability more strongly than complexity, warming or invasion

**DOI:** 10.1101/2025.01.27.634676

**Authors:** Miguel G. Ximenez-Embun, Gregoire Proudhom, Mélanie Thierry, Nicholas A. Pardikes, Anna Mácová, Jan Hrček

## Abstract

Ecological communities face major challenges of biodiversity loss, invasive species, and warming due to human action. Factors that determine the dynamics, stability and functioning of network of interacting species have been debated in ecology since its beginning as discipline. Community complexity and dominant (keystone) species have been proposed as the most relevant factors, however there is lack of knowledge on how these factors will interact with warming and invasion. To determine the effect of complexity, warming or invasion and their combinations on stability and functioning of a food web we performed a microcosm laboratory experiment using *Drosophila*-parasitoid food webs as a model. During five months, we recorded the dynamics of twelve different food webs (different combinations of flies and parasitoid species with two complexity levels) subjected to two temperature regimes (ambient and warming) and invasion or not by *Drosophila suzukii* in a factorial design. We did not observe any major effect of complexity or invasion. However, species composition had major effect, as webs with *Drosophila simulans* present were rapidly dominated by it, reducing diversity, increasing parasitism rate, reducing stability and bringing the community on the verge of collapse. Warming in turn increased diversity, overall fly population and temporal stability, while decreasing parasitism rate. Invasion of *D. suzukii* was not successful in any of the networks, possibly due to its low competitive ability. Our results imply that dominant species can have very strong effects on dynamics and stability, compared to food web complexity.

## Introduction

Most ecological communities face strong pressure from human population, with biodiversity loss, warming and invasive species being among the most prominent challenges for community stability and functioning (Hooper et al., 2005; Tylianakis & Binzer, 2014; Wardle et al., 2011). The role of diversity has been widely studied since the beginning of ecology as a discipline (Eschenbrenner & Thébault, 2023; Tilman & Downing, 1994). However, the effects of warming, invasion, and combined challenges on community dynamics and functioning are much less understood and need more research to help us understand and mitigate adverse effects. Especially so in communities with strong feedbacks between trophic levels, as community stability studies are commonly restricted to a single trophic level.

### Effect of complexity on food web stability and functioning

Researchers have for long time been interested in factors determining stability and functioning of networks of interacting species. By far the most studied factor has been the diversity or complexity of the system. More complex ecological communities (with higher number of species and higher number of interactions) are understood provide more ecosystem functions, to be more temporary stable, more resilient to invasive species (Levine & D’Antonio, 1999) and to extinctions (Steiner et al., 2005; Xu et al., 2021). However, there are also many exceptions (Ives et al., 2000; Sasaki & Lauenroth, 2011; Zhao et al., 2023). On the other hand, we know that some species can have disproportionately strong effect and largely determine community stability and functioning (Bowker et al., 2021; Sasaki & Lauenroth, 2011; Segrestin et al., 2024). These species have been variously called keystone, or dominant, and their presence can completely transform the system (Segrestin et al., 2024; Tilman & Downing, 1994), like in the famous sea star example (Menge et al., 1994). Invasive species are also known to often have disproportionate effects (David et al., 2017). Major advantage of experimental studies is that they can separate the effects of complexity and dominant species (Wardle, 1999).

In food webs, fluctuations due to overexploitation of resource species by consumers are natural part of the dynamics (Hiltunen et al., 2014). To understand temporal stability and resilience it is therefore important to follow the dynamics, not only its end point or average (Fox, 2007; Tylianakis et al., 2006). Very strong fluctuations increase the probability of extinctions (Pimm et al., 1988). On the other hand, too weak fluctuations may signal loss of top-down control function which can lead to cascading extinctions through release of competitively strong species (Pardikes et al., 2022). Top-down control regulates population dynamics and has a crucial role in ecosystem stability (Pedroso et al., 2021; Tylianakis & Binzer, 2014). Stability of fluctuating systems is still little understood, but presence of a combination of strong (dominant) and weak links is expected to enhance persistence (O’Gorman & Emmerson, 2009). Changes in one trophic level may have consequences in others (Rezende et al., 2021). For instance, diversity of predators had a negative effect on prey temporal stability (Halpern et al., 2005), while diversity of parasitoids enhanced it (Tylianakis et al., 2006) or had neutral effect (Rodríguez & Hawkins, 2000).

### Effect of warming and invasions on food webs

Warming changes food web dynamics, altering individual species performance and their interactions. However, the effects are not very clear and are community specific. Warming in communities with differential thermal tolerance among species can lead to changes in species relative abundances and dominance at one trophic level (Olsen et al., 2016) or between trophic levels (Petchey et al., 1999; Rezende et al., 2021; Synodinos et al., 2021). It alters community functioning by increasing predation rate (Vucic-Pestic et al., 2011; Walker et al., 2020) or decreasing parasitism rate (Thierry et al., 2019; Tylianakis & Binzer, 2014), but the consequences for resource species abundances is not consistent (Thierry et al., 2019; Vucic-Pestic et al., 2011). The effect of warming on the resource-consumer temporal stability is not consistent; it has been shown to be negative (Zhao et al., 2023), neutral (Rezende et al., 2021), or positive (Gårdmark & Huss, 2020; Vucic-Pestic et al., 2011). Diversity in aquatic communities might buffer the effect of temperature on stability (Petchey et al., 1999) but is not always the case (Zhao et al., 2023). There are few studies on diversity and warming for terrestrial ecosystems.

Invasion also affects food web dynamics and stability. The effect depends on the trophic level invaded. Invasions at consumer level tend to have a positive bottom-up effect increasing food web stability while invasive predator might have a destabilizing effect (David et al., 2017; Duan et al., 2024). The factors that determine the invasibility of a community are still controversial (Hui & Richardson, 2019), but more diverse communities are expected to be more resistant to invasion (Levine & D’Antonio, 1999). Invasion is also used as a proxy to study the resilience of food webs by examining recovery from the invasion (Donohue et al., 2016). Warming can increase the susceptibility of food webs to invasions, reducing the community stability (Sentis et al., 2021).

### Objectives and study system

In this study, we used a microcosm experiment to test the effects of complexity, dominant species, warming, invasion and their combinations on stability and functioning of a terrestrial food web. Microcosm experiments following the dynamics of food webs are very challenging but offer unique opportunity to separate the effect of the different factors (Petchey et al., 1999; Steiner et al., 2005). As model system we used *Drosophila*-parasitoid food web from tropical Australia (Jeffs et al., 2021). As invasive species we chose *Drosophila suzukii* because it is the only Drosophilid fly that is considered a pest and invasive species and is causing big economic losses (De Ros, 2024; Tait et al., 2021). We assembled food webs with two complexity levels while accounting for the effect of individual species. We then exposed the webs to ambient or warming temperatures and invasion of *D. suzukii* and studied resulting dynamics for five months. We analysed: (1) species richness and diversity and their dynamics, (2) the overall community population dynamics and its top-down control function (parasitism rate) and (3) temporal stability of the mentioned parameters.

## Materials and methods

### Study system

We used a community of *Drosophila* flies and their parasitoids from Australian rainforest as a study system. We used eight species of flies: *Drosophila simulans (SIM), D. pesudotakahashii (PST), D. bipectinata (BIP), D. pseudoananassae (PSA), D. birchii (BIR), D. bunanda (BUN), D. pallidifrons (PAL), D. sulfurigaster (SUL)*. We collected all lines between 2017 and 2018 at two altitudinal gradients in North Queensland Australia: Paluma (S18° 59.031’ E146° 14.096’) and Kirama Range (S18° 12.134’ E145° 53.102’), between 70 and 880 m.a.s.l. (Jeffs et al., 2021). We created isofemale lines, identified them by morphology and DNA barcoding, and shipped them to Czech Republic under permit no. PWS2016-AU-002018 from Australian Government, Department of the Environment. Lines were maintained at 23 ºC and 12/12 hours light:dark regime on standard *Drosophila* medium (corn flour, yeast, sugar, agar and methyl-4-hydroxybenzoate) for approximately 45 to 70 non-overlapping generations before the experiment. To recover genetic diversity, 2-8 isofemale lines of each species were mixed into a mass bred line five generations before the experiment, when possible, we included isofemale lines from the two sites and higher and lower elevations. We used genetically variable culture of *D. suzukii* as invader species, collected in Trentino (Italy), established in the laboratory by Gianfranco Anfora team from Fundazione Mach, and shipped to Czech Republic.

We used four currently undescribed species of hymenopteran parasitoids from the same Australian rainforest food web. One pupal parasitoid, *Trichopria* sp. (Diapriidae; strain 66LD reference voucher no. USNMENT01557254, reference sequence BOLD process ID: DROP096-21) and three larval parasitoids: *Ganaspis* sp. (Figitidae; strain 75B reference voucher no. USNMENT01557101, reference sequence BOLD process ID: DROP164-21), *Leptopilina* sp. (Figitidae; strain 106BPL reference voucher no. USNMENT01557104, reference sequence BOLD process ID: DROP053-21) and *Asobara* sp. (Braconidae; strain KHB, reference voucher no. USNMENT01557097, reference sequence BOLD process ID: DROP043-21). For more details on the parasitoid strains see Lue et al. 2021. *Ganaspis* sp. (G) and *Trichopia* sp. (T) were able to develop in all fly species while and *Asobara* sp. (A) in all but BIP, PSA and PST and *Leptopilina* sp. (L) in all but BIP PSA, PST and SUL. We maintained parasitoid species for approximately 25 to 40 non-overlapping generations before the experiment on *Drosophila melanogaster*, so that they do not adapt to one of the species from the experimental community.

### Design of the communities

We constructed food webs of two levels of complexity: high (4 flies x 3 parasitoid species) and low (3 flies x 2 parasitoid species) from mass bred lines of individual species. According to previous data 10 interactions are expected on high complexity webs while 5 in low complexity ones (2-fold difference). We standardized connectance at 0.83, to not confound the effect of complexity with connectance. Each level of complexity was replicated six times with different species composition. One of the main constrains of interpreting biodiversity experiments is sampling effect, where presence of particular species is confounded with the effect of diversity or complexity *per se* (Huston, 1997; Wardle, 1999). To avoid this confounding effect, we ensured that every species is present in the same number of replicates of high and low complexity. Species composition of each food web is presented in Table 1. We set and maintained each food web in a plastic box (47cm x 30cm x 27.5cm) with holes on sides covered with mesh tissue for ventilation.

**Table 1.** Species composition of the twelve food web (six high and six low complexity). For species abbreviations see beginning of Materials and Methods section. Temporal blocks were as follows: Block 1: H1, H4, L1, L4. Block 2: H2, H5, L2, L5. Block 3: H3, H6, L3, L6.

| HIGH complexity (4 Flies x 3 Parasitoids) |  |  |  |  |  |  | LOW complexity (3 Flies x 2 parasitoids) |  |  |  |  |  |  |
| --- | --- | --- | --- | --- | --- | --- | --- | --- | --- | --- | --- | --- | --- |
| web | Flies |  |  |  | Parasitoids |  |  | web | Flies |  |  | Parasitoids |  |
| H1 | SIM | BUN | SUL | PST | A | L | G | L1 | SIM | SUL | PST | A | G |
| H2 | SIM | BUN | PAL | PSA | A | L | T | L2 | SIM | BUN | PSA | L | T |
| H3 | BIR | BUN | SUL | PSA | A | L | T | L3 | SUL | BUN | PSA | L | T |
| H4 | BIR | PAL | BIP | PSA | L | G | T | L4 | BIR | PAL | BIP | L | T |
| H5 | BIR | SUL | BIP | PST | A | G | T | L5 | BIR | SUL | BIP | A | G |
| H6 | SIM | PAL | BIP | PST | A | G | T | L6 | SIM | PAL | PST | A | G |
|  | universal host |  |  |  | universal parasitoid |  |  |  |  |  |  |  |  |
|  | host of A, T or G |  |  |  | non universal parasitoid |  |  |  |  |  |  |  |  |
|  | only host of T or G |  |  |  |  |  |  |  |  |  |  |  |  |

### The experimental design

In addition to web complexity factor introduced above, the experiment involved warming and invasion (introduction of *D. suzukii*) factors. The design was fully factorial, including all combinations of complexity, warming and invasion for a total of 48 food webs (boxes). Each of the twelve webs (six with low and six with high complexity) was subjected to i) ambient temperature of 23ºC, ii) warming temperature of 27ºC, iii) invasion at ambient temperature, and iv) invasion at warming temperature. Ambient temperature corresponds to current average temperature in the Australian rainforest understory where the community is coming from. In a climate change model, a 1-6 °C increase in temperatures by 2070 is predicted (IPCC 2014), thus 27°C represents a general warming scenario in tropical Queensland, Australia. The experiment lasted six months. In the invasion treatments, we introduced *D. suzukii* on day 63 (approximately two months), after the last introduction of larval parasitoids. We chose *D. suzukii* due to its importance as invasive species in Europe and North America (De Ros, 2024; Tait et al., 2021). *Drosophila suzukii* has so far not been recorded from Australia and we therefore tested its ability to invade the studied rainforest community. The experiment was carried out in growing chambers with controlled conditions with a L:D 12:12 photoperiod and 65% relative humidity. During the experiment some boxes had to be eliminated due to the appearance of mould. At the end of the experiment, they were 34 boxes left from the initial 48.

### Food web establishment and invasive species introduction

We divided the food webs designs into three temporal blocks of four food webs each, offset by two days to make running of the experiment feasible (see Table 1). To create as asynchronous food webs as possible, we introduced flies in five consecutive dates every two days, introducing 12 adults (half male, half female) per species per date (a total of 60 adults/species per cage). That meant initial population of 180 adult flies/cage in low complexity food webs and 240 flies/cage in high complexity webs. We introduced parasitoids following the same scheme but introduced larval parasitoids two days after flies and pupal parasitoid (T) 8 days after flies, to allow development of host life stage. We imposed an initial parasitism rate of 25% comparable to natural food webs (Jeffs et al., 2021). We thus introduced four adults (two male, two female) per time per species, in total 44 parasitoids/cage in low complexity and 60 parasitoids/cage in high complexity.

We introduced invasive species 63 days after last parasitoid introduction and in the same way as the other fly species (five times, two days apart). We set the invasion ratio at 10% of average of pupae present in sampling of the first time point (day 32). The invasion ratio is similar to previous studies (Hausch et al., 2018). On each introduction, 18 fly adults (half male, half female) were introduced in low complexity boxes and 24 fly adults into high complexity boxes, for a total of 90/120 introduced flies per box.

### Experiment maintenance and sampling

We introduced standard glass *Drosophila* vials with 10 ml of diet (same diet as described above) to the boxes to allow flies to lay eggs and develop. We introduced new vials every five days, and each vial was inside the box for 40 days, enough time for the species with the slowest development time to emerge (*Ganaspis* sp., 30-35 days). In high complexity boxes, we introduced five vials each time, for a maximum total of 40 vials in a box. In low complexity boxes we introduced four vials (and only three vials once every twenty days) to reach a total number of 30 vials in the box. In that way a ratio of approximately 0.75 of food vials, as well as fly and parasitoids initial population was maintained between low and high complexity levels.

### Sampling, rearing and species identification

We sampled the webs for species composition and population dynamics 1, 3, 4 and 5 months after the last introduction of larval parasitoid. We did the first sampling before invasion and the rest after invasion (of the invasion treatment webs). We did two sub-samplings at each sampling time with four days apart to reduce the chance of missing rare species and species with shorter development time. On each sub-sampling, we introduced 2/3 vials in low/high complexity boxes (that represent 15% of the total vials on the box) and left them in the box for nine (in boxes at 27 ºC) or eleven days (at 23 ºC). That time was enough for the flies to pupate and be exposed to pupal parasitoids. We spread pupae from sampling vials on a tray, photographed them to be later counted using ImageJ software (Schindelin et al., 2012) and randomly selected 192 pupae (by taking pupae closes to pre-made dots on the tray), placed in 96-wells plates and reared in growing chambers (23ºC or 27 ºC depending on the temperature treatment, 65% RH) till the emergence of flies or pupae. We identified the pupae and emerged adults of flies or parasitoids mostly morphologically but when it was not possible then molecularly by sequencing the COI locus.

### Population dynamics and parasitism rate

We calculated the ratio of emerged adults of every species out of the 192 sampled pupae. That ratio was multiplied by the total number of pupae in the sampling vials and divided by the ml of food introduced in the sampling vials (40/60 ml in low/high complexity treatment) to get number of emerged adult per ml of food. Parasitism rate was calculated as the number of parasitoids emerged divided by the sum of flies and parasitoids emerged.

### Measuring diversity and stability

We evaluated the effect of treatments on species richness, Shannon diversity and Pielou’s index of evenness at each timepoint, as we expected a fluctuating dynamics affected by the treatments. Richness is simply the number of species emerged. We chose Shannon index (H’) due to the big imbalance of relative abundance between species. It was calculated as:

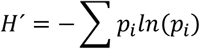

Where pi is the relative abundance of every species calculated as the fraction of the adults emerged of one species divided by all total emerged adults on that box. As indicator of dominance we used Pielou’s index of evenness calculated as the ratio between H’ and the logarithm of the species richness.

To measure temporal stability of adults emerged and of Shannon diversity index, we used inverse of covariance, ICV=μ/σ. Where μ is the average of relative abundance of emerged adults of every species in a box over time, or average of Shannon index of every box over time while σ is the standard deviation of the same metric. ICV is commonly used to measure temporal stability (Sasaki & Lauenroth, 2011; Tsafack et al., 2019).

### Statistical analysis

We used warming, complexity and invasion as predictors and every food web design was considered as a replicate within each complexity level (6 replicates in low and 6 in high complexity). As frequency of all species was balanced, we did not have an *a priori* expectation on individual species performance or whether any species will influence the dynamics of the food webs the most. We expected that no species will stand out at all levels of temperature, initial complexity and invasion treatments. However, after visually examining the dynamics of all food webs (Figure S1) we saw that *D. simulans* strongly dominated the food webs where it was present, often reaching more than 95% of the total population (when present). We therefore decided to add presence of *D. simulans* as additional predictor including its interaction with warming. This is possible, because *D. simulans* was present in half of the initial food webs.

We did the analysis separately for every sampling time point (month 1, 3, 4 and 5). We analysed the data using generalized linear models with appropriate distributions. We analyzed response variables Richness, Shannon index, Pielou’s eveness and ICV using normal distribution after testing for normality and log-transforming when needed. For Parasitism rate we used a quasi-binomial distribution and for fly and parasitoid emergence we used quasi-poisson distribution. We did all analyses using R 4.1.2 (Team, 2021).

## Results

### Diversity

Warming and presence of *D. simulans* were the most important factors determining the diversity indexes measured and their dynamics. The effects depended on the parameter measured and the time point because of fluctuating dynamics.

Warming had a significant effect on the diversity indexes that include the relative abundances of species: diversity and evenness. However, the effect was visible only at the end of the experiment. At month four, the warming treatments had significantly higher diversity and evenness (Table 2, Figure 1). At month five this trend was maintained in the food webs with *D. simulans* but was lost in food webs without it.

**Table 2.** Statistics (t-value) of the main model predictors for species richness, Shannon diversity and evenness. The code m1-5 signifies the month of the sampling. The colours represent the level of significance (see legend below the table).

|  | Richness |  |  |  | Shannon diversity |  |  |  | Evenness |  |  |  |
| --- | --- | --- | --- | --- | --- | --- | --- | --- | --- | --- | --- | --- |
|  | m1 | m3 | m4 | m5 | m1 | m3 | m4 | m5 | m1 | m3 | m4 | m5 |
| **Warming** | -1.2 | -1.4 | 0.3 | 0.8 | -0.7 | 0.6 | 2.3 | -1.3 | -0.3 | 1.7 | 2.3 | -1.7 |
| **Initial complexity** | -2.3 | -4.1 | -4.1 | -3.7 | -1.4 | -1.6 | -1.6 | -0.4 | -0.3 | 1.0 | 1.2 | 1.9 |
| **Invasion** | -0.2 | 0.2 | 0.2 | 0.9 | -0.3 | 1.2 | 0.3 | -0.2 | 0.0 | 1.1 | 0.2 | -0.4 |
| ***D. simulans*** | -1.9 | -3.3 | -3.3 | -1.2 | -4.1 | -4.4 | -5.6 | -4.3 | -3.8 | -2.0 | -4.5 | -4.4 |
| ***W27°C:D.sim*** | -1.0 | 0.7 | -0.6 | 0.0 | 1.1 | -1.5 | 0.1 | 2.8 | 1.3 | -2.7 | 0.5 | 3.7 |
| **p-value** |  | < 0.05 |  | 0.01 |  | <0.001 |  |  |  |  |  |  |

**Figure 1.**
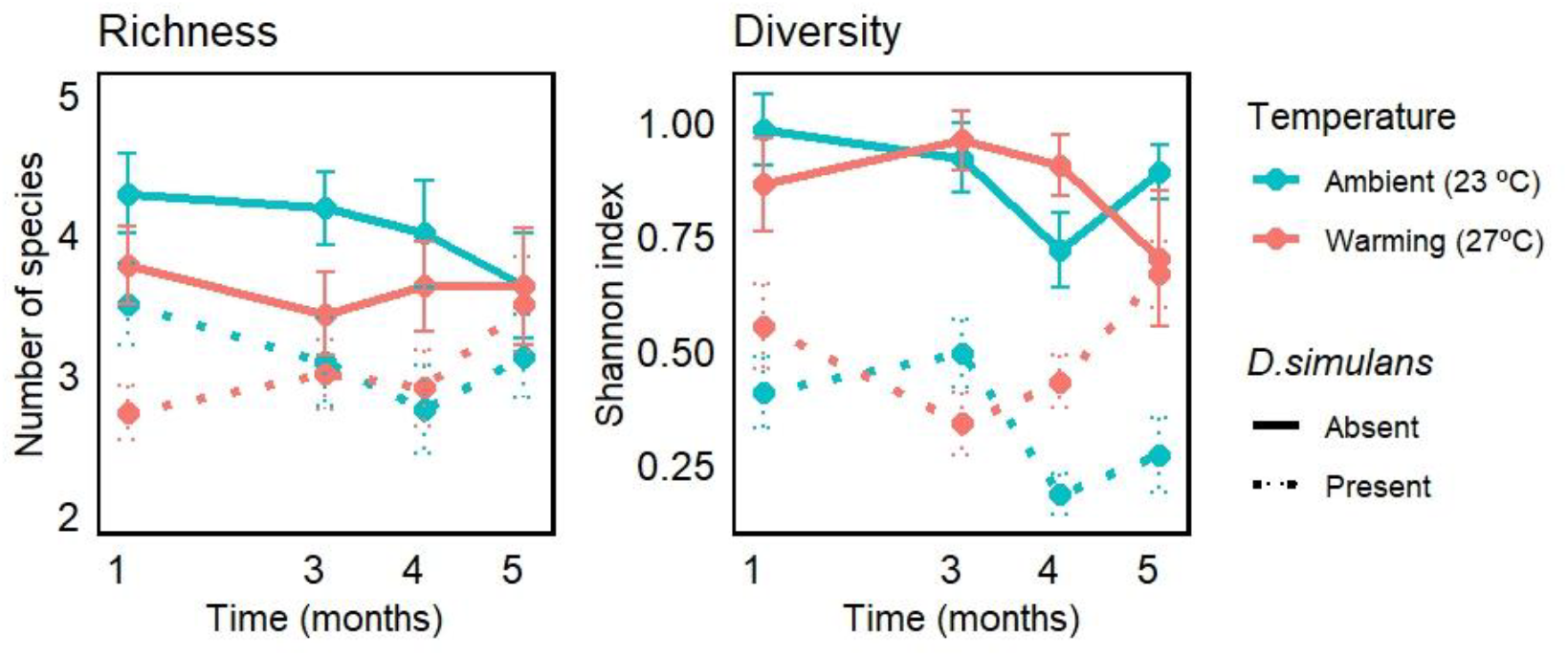
Dynamics of species richness and diversity depending of temperature regime and presence/absence of *D. simulans*.

The presence of *D. simulans* in the food web had a major effect on all aspects of the dynamics during the entire experiment (Table 2, Figure 1). It significantly reduced species richness in months three and four, and strongly reduced diversity and evenness at all timepoints.

Food webs with higher initial complexity had significantly higher species richness according to expectations, and this effect was maintained throughout all timepoints (Figure 1 and Table 2). Interestingly, initial complexity did not affect diversity or evenness during the experiment. There was a decrease of about two species from the initial richness of five or seven species (low and high complexity treatments) at month one, followed by more or less stable richness during the rest of the experiment (Figure 1). As consequence, there was no effect of other treatments on final richness (Table 2).

Invasion by *D. suzukii* which took place just after sampling in month one did not have a significant effect on species richness, diversity or eveness (Table 2). The invasion of *D. suzukii* was not successful in any of the treatments and replicates.

### Population dynamics and parasitism rate

We observed fluctuating host – parasitoid dynamics with initial decrease in fly numbers, followed by an increase and corresponding changes in parasitoid emergence and parasitism rate (Figure 2; for dynamics of individual replicates see Figure S1). Like for diversity, warming and the presence of *D. simulans* were the factors with significant effect on emergence dynamics and parasitism rate.

**Figure 2.**
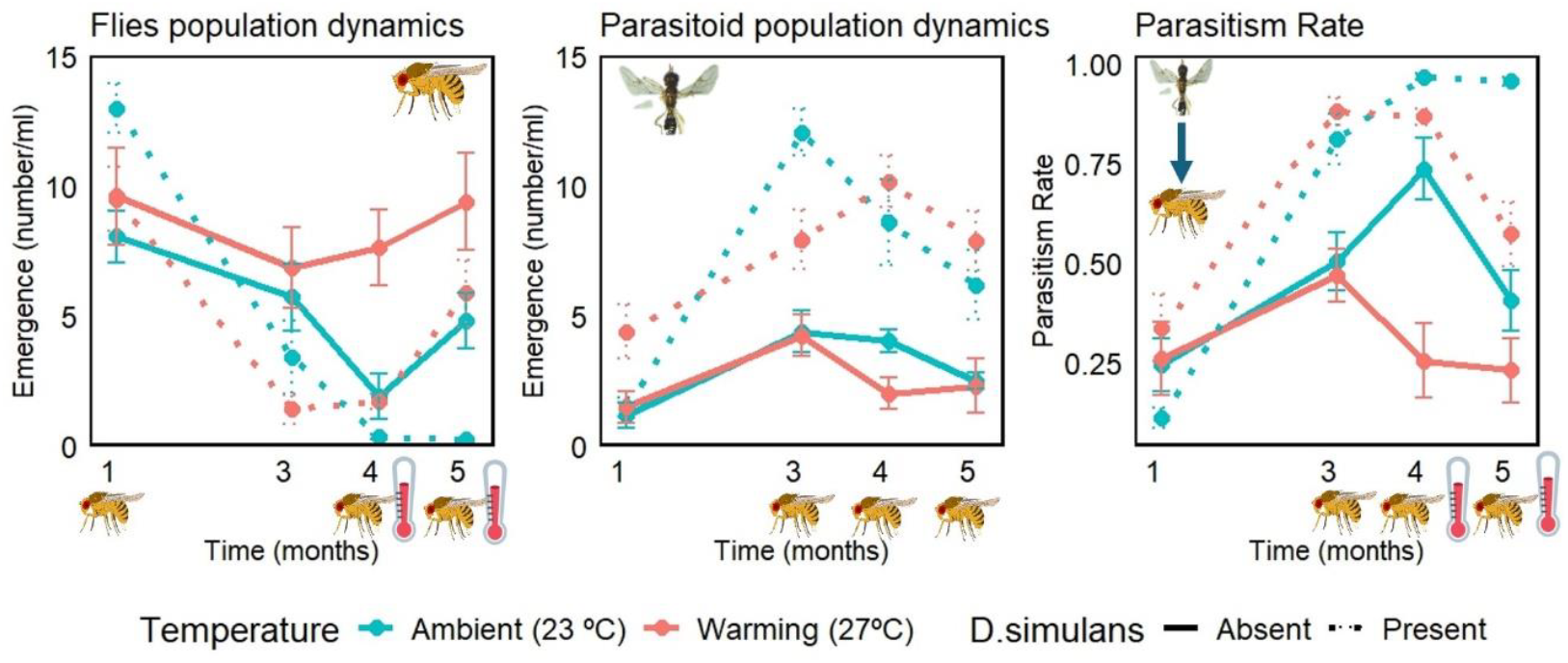
Population dynamics of emerged adults (flies and parasitoids) and parasitic rate depending on warming and presence of *D. simulans*. Thermometer and fly icons represent treatments that showed a significant effect of warming or *D. simulans* presence at a given timepoint (see Table S1).

The effect of warming was significant in months four and five, where fly emergence when subjected to warming was higher and parasitism rate was lower (Figure 2 and Table S1). Parasitoid emergence was not significantly affected by temperature.

The presence of *D. simulans* had a positive significant effect on parasitoid emergence and parasitism rate from month three onward. The effect on fly emergence was more complex, with a significant positive effect in month one, which turned into a significant negative effect at months four and five. The fly emergence in presence of *D. simulans* got very low at the end of the experiment, particularly in the ambient temperature treatment, while parasitoid emergence and parasitism rate were very high (Figure 2, Table S1).

### Temporal stability

The magnitude of natural occurring fluctuations is commonly used a measure of the stability of the system. In this case we used the inverse of covariance which indicates more temporal stability.

Warming significantly increased the temporal stability of total fly emergence but not total parasitoid emergence. In contrast, the presence of *D. simulans* decreased the stability of fly and parasitoid emergence, and also community diversity and eveness (Figure 3 and Table S2). None of the factors influenced inverse of covariance of parasitism rate (Table S2).

**Figure 3.**
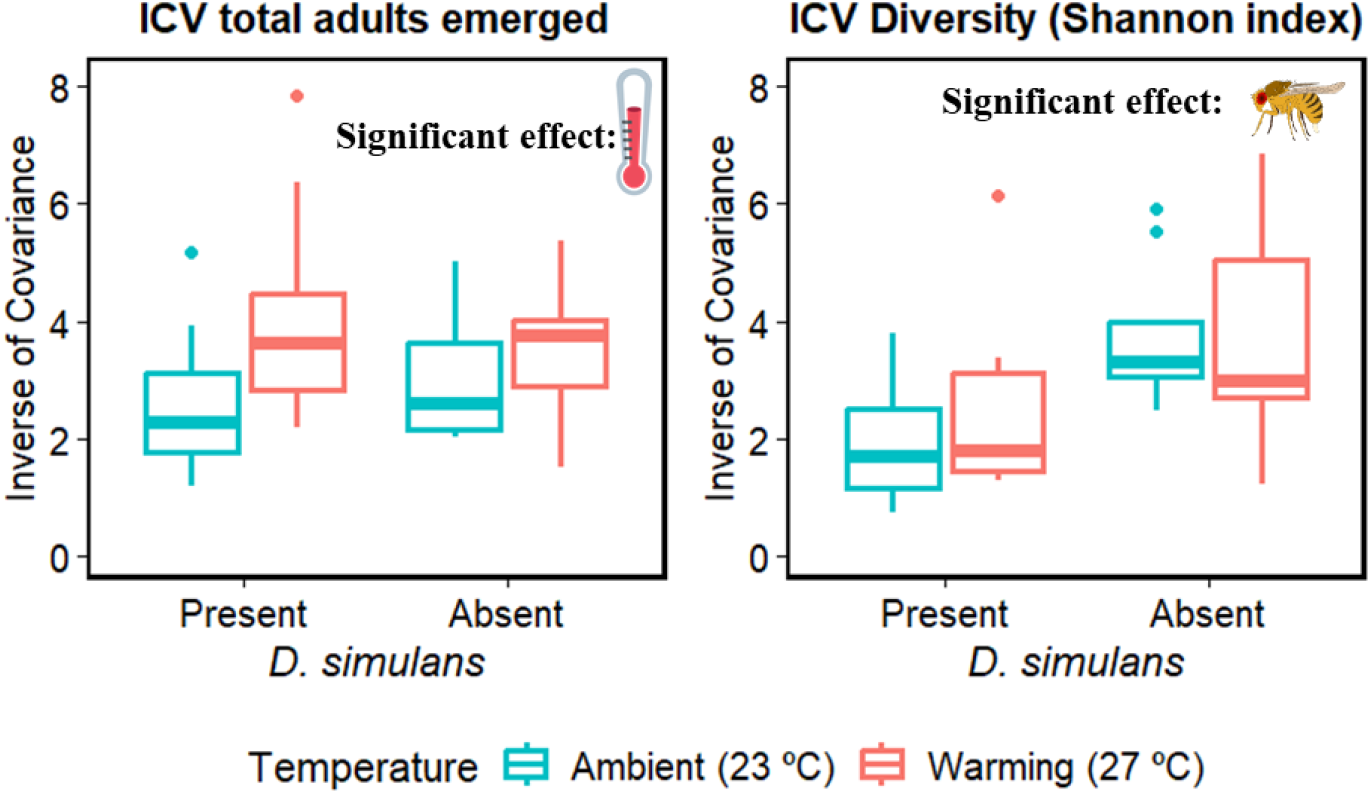
Temporal stability (Inverse of covariance) of emerged fly abundance and Shannon diversity. Thermometer and fly icons represent treatments that showed a significant effect on ICV of warming or *D. simulans* presence respectivelly (see Table S2 for statistics).

## Discussion

We performed a long-term microcosm experiment to explore whether complexity, warming, invasion or species composition determine food web dynamics and stability. We found unexpectedly low effects of 4°C warming, major step in initial complexity (1.4-fold species richness, 2-fold potential interactions between complexity levels) and invasion of an important pest. Instead, species composition had major effect. When *Drosophila simulans* was present, it rapidly dominated over other *Drosophila* flies reducing diversity, increasing parasitism rate, reducing stability and bringing the community close to collapse. Warming in turn showed effect on diversity and parasitism rate only after 3 months and increased diversity, overall fly population and temporal stability, while decreasing parasitism rate.

### Effects of dominant species (*D. simulans*) on food web stability and dynamics

The factors that determine food web stability and dynamics have been long debated. The species-specific effect on communities has been a concern for long time in biodiversity ecosystem function studies as they can confound the biodiversity effect by sampling (Huston, 1997; Wardle, 1999). At the same time, later studies state that is not only the richness of species what matters but also the relative abundance of the species present (Bowker et al., 2021). In this regard, plant competition studies observed that in communities with high dominance of one species, the dynamics, stability and functioning is more determined by that species, than by complexity per se (Sasaki & Lauenroth, 2011; Segrestin et al., 2024). However, this effect was hardly ever shown in multitrophic terrestrial system. We observed that *D. simulans* dominated all food webs when present. The dominance by one species in drosophilid communities has been observed before in experiments with three fly and one parasitoid species (Davis et al., 1998), and in nature in temperate areas with low diversity (Galludo et al., 2020). The differences in competitive ability among *Drosophila* species might explain the dominance as stated by coexistence theory that predicts the superior competitive ability of *D. simulans* or *D. sulfurigaster* and the low competitive ability of *D. pseudotakahashi* (Terry et al., 2021) and *D. birchii* (Chen & Lewis, 2023). *D. simulans* excluded the other flies species at the two middle samplings (month 3 and 4) with a significant reduction in species richness (Table 2), but there was some diversity recovery at the end (Figure 1). This means that some species fell below detection power of the sampling at months 3 and 4. *D. similans* induced an increase in parasitism rate, increasing parasitoid emergence (Figure 2). That led to reduction of fly emergence to a level that the webs could have collapsed if we were able to follow them for longer (Pimm et al., 1988). Alternatively, the fly population could recover and the community could enter a cyclical dynamic. Longer experiment would show which of the two scenarios would take place.

*D. simulans* decreased temporal stability of both fly and parasitoid trophic levels (Table S2), but the effect was stronger on the lower trophic level (flies) than on the parasitoids as previously observed in other studies (Jiang & Pu, 2009; Rezende et al., 2021; Zhao et al., 2023).

### Effects of complexity

Our experimental design allowed testing the effect of initial complexity, while standardizing initial connectance and correcting for sampling effect. The difference between high and low complexity treatments (1.4-fold in species richness, 2-fold in potential interactions between complexity levels) was high compared to natural differences between food webs, for instance in Jeffs et al. (2021). Despite that, there was very little effect of complexity. One possible explanation is that while the difference in richness between complexity levels was maintained (Figure 1) there was not observed difference on number of interactions or connectance at any time point. Experimentally, it would also not be possible to create a bigger difference and at the same time keep controlling for sampling effect and standardizing connectance. However, it is possible that complexity in nature can act through connectance, it has shown to be a key factor for food web stability (Thébault & Fontaine, 2010), exploring this would require additional experiments.

### Effects of temperature and invasion

Warming can have variable effects on stability (Gårdmark & Huss, 2020; Rezende et al., 2021; Zhao et al., 2023). In our case it had a stabilising effect, significantly reducing the fluctuations in overall abundance (Figure 3). In case of the webs with *D. simulans*, warming kept the fly population at a level that might prevent the collapse that was likely at ambient temperature (Figure 2). The effect of warming on stability can be explained by two main mechanisms: the effect of warming on individual species (Gårdmark & Huss, 2020) or the effect on the trophic interactions (Vucic-Pestic et al., 2011). Warming has differential effect on the *Drosophila* species from our study system (Chen & Lewis, 2024). However, we did not observe warming favouring some species over others or even increased dominance as has been shown by a single generation experiment (Thierry M et al., 2021). This was probably because the three most dominant species (*D. simulans, D. bipectinata* and *D. sulfurigaster*) were close to their thermal optimum in both ambient and warming treatments (Chen & Lewis, 2024). Warming also increased overall fly emergence (Figure 2). This effect could be a combination of better performance of the dominant fly in warming or the reduction in parasitism rate. Indeed, warming influenced parasitoids performance, reducing parasitism rate as previously described (Thierry et al., 2022). However, we did not observe effect of warming on overall parasitoid emergence (Figure 2), as the increase of fly population by warming compensated the reduction in parasitism rate. Warming only increased temporal stability in overall fly emergence and not in parasitoid emergence (Table S2). In prey-predator dynamics a reduction in predator efficiency induced by warming (in our case reduction of parasitism rate) dampens oscillations and increases temporal stability (Rall et al., 2010; Vucic-Pestic et al., 2011). We thus suggest that the increase in temporal stability of overall fly emergence is due to the reduction of parasitism rate.

*Drosophila suzukii* failed to invade all experimental food webs and we did not observe any pupae or emerged flies in our samples despite relatively high invasion ratio of 10% of the fly community. Possible reason for this is that *D. suzukii* is a poor competitor (Shaw et al., 2018). It is the only drosophilid fly that can pierce the skin of a fruit and can thus be the first drosophilid to invade the fruit patch. Competition has been shown to have stronger effect on *D. suzukii* than parasitism (Dancau et al., 2017). Failure to invade in a wide variety of complexity and thermal treatments as well as 6 different species compositions demonstrated here presents a very strong evidence that *D. suzukii* cannot withstand competition by other species. However, lack of invasion success should not be taken as evidence that *D. suzukii* cannot invade the Australian rainforest community in nature, as the food (fruit) availability and other conditions will be very different form our microcosm settings. Based on its establishment in Europe and North America we expect *D. suzukii* would be able to invade Australian communities if it was introduced there.

## Conclusions and future directions

Our study shows that effects of dominant keystone species should not be neglected as they might determine the dynamics and functioning of a web. This should be particularly considered when designing Biodiversity Ecosystem Function experiments as previously stated (Wardle, 1999). Our study points the importance of dominance in a multitrophic community, where we observed that the dominant species determines the dynamics of communities and how are they affected by perturbations like warming. To understand the mechanisms and food webs functioning, it is important to examine the dynamics at all trophic levels. Finally, although we have seen a stabilising effect of warming on all the mentioned processes there is still a big gap of knowledge on how the combination of changes in complexity, warming and invasion affect food webs dynamics that should be further explored.

## Acknowledgements

We thank Martin Libra, Joel Brown, Andrea Weberová, Inga Freiberga, Anna Jandová, Shafia Zhara, Michaela Borovanská and Eva Chocholová for their help with the pupae sampling. We thank Eva Kriegová, Varvara Fedorchenko, Kateřina Lávičková and Tereza Volfová for help with identification of the insects. The study was funded by the Czech Science Foundation grant number 20-30690S. MGX-E was additionally supported by “European fellowships H2020 - LeishOmics and Invaweb” grant no. CZ.02.2.69/0.0/0.0/19_074/0016248 from Czech Ministry of Education, Youth and Sport, co-funded by European Union.

## Authors’ contributions

MGXE and JH conceived the project; MT, GP and NAP contributed to the experimental design; MGXE, MT, NAP, GP and AM collected the data; MGXE and JH analyzed the data. MGXE and JH wrote the manuscript and all authors contributed critically to the drafts and gave final approval for publication.

## Data Availability Statement

All raw data and R scripts used for this study will be made available through Zenodo.

