## Supplement Material for "Dominant species determines *Drosophila*-parasitoid food web dynamics and stability more strongly than complexity, warming or invasion"

### SUPLEMENTARY MATHERIAL

**Rules for foodweb design.**

- Same initial potential structure all the webs (expected initial conectance 0.83)
- 4x3 webs have double of possible trophic interactions (10) than 3x2 webs (5).
- 4x3 webs half with:
  - 2 generalist parasitoids + 1 "specialist" AND 2 host attacked by all parasitoids 2 attacked only by generalist
  - with 1 generalist parasitoids + 2 "specialist" AND 3 host attacked by all parasitoids 1 attacked only by generalist
- All the 4X3 webs differ al least on 2 host species among them
- The 4x3 webs we have 4 different combinations of parasitoids the two repeated ones are coupled with host combinations that have only one host in common.
- The host of 3x2 webs are selected to try to copy the same interaction among host than the 4x3 (the webs 1-3 have a similar version in the 3x2 taking out one species)
- Regarding to bloc:
  - On each bloc in 4x3 one time each species.
  - On the 3x2 bloc we make similar combinations than in 4x3 and put them in same bloc.
- Host combinations: Try to avoid to have two species from the same subgroup n the same web: SIM-PST, SUL-PAL, BUN-BIR, PSA-BIP as that will make more difficult the identification
- Host parasitoids interactions (we avoid the combinations were the parasitoid control is loosed):
  - Try to avoid the combination of BIP with two of G, L or A
  - T, better PSA or BIP than PST
  - SUL better controlled by G and A
  - PAL better T and L

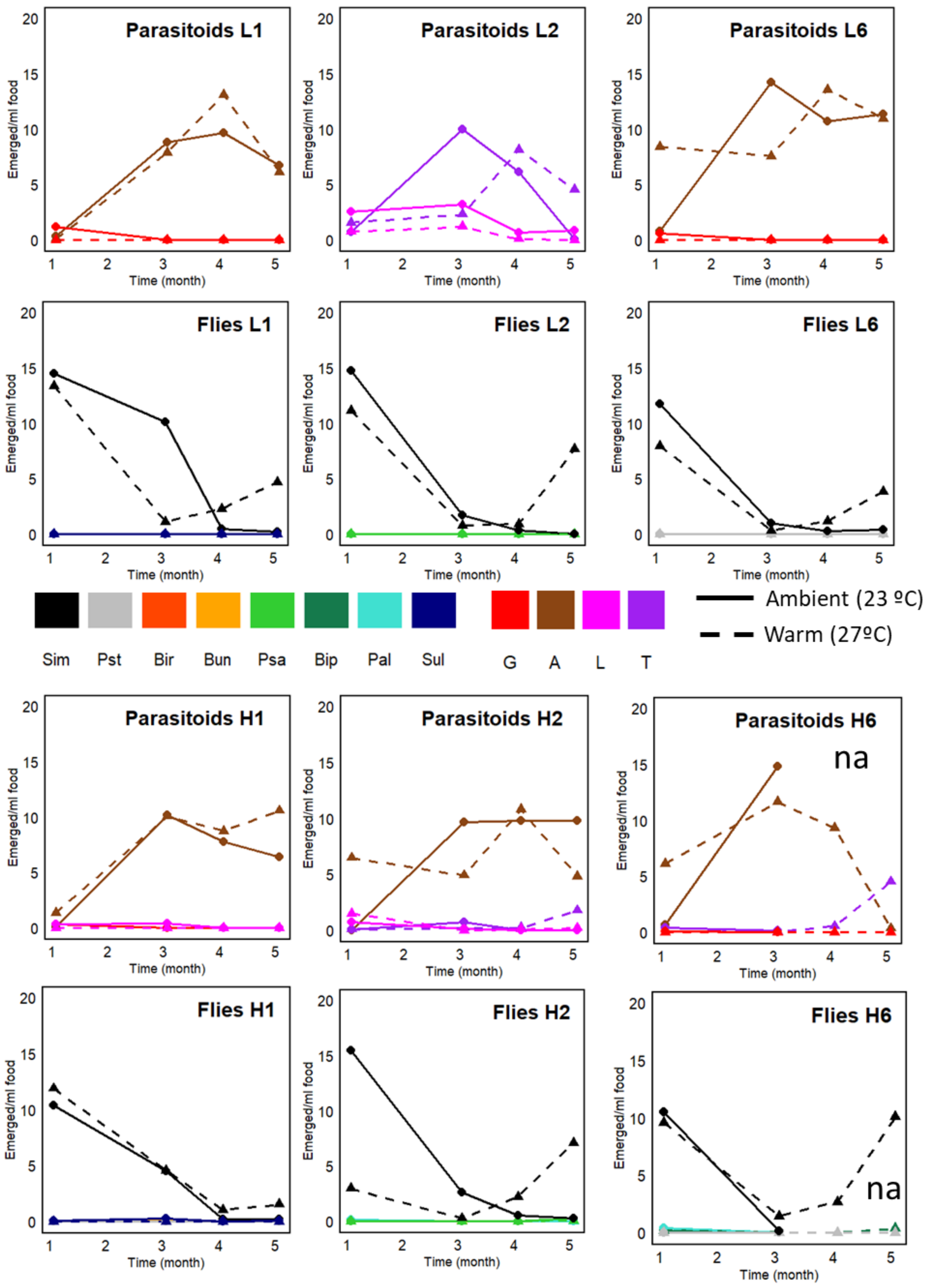


**Suplementary figure 1A.** Population dynamics (number of emerged adults/ml of food) of the different species in the food webs designs (L: low, H: high complexity) with presence of *D. simulans* during 5 months. Solid line represent the boxes under ambient temperature (23 °C) and dashed line under warming (27°C). The data show the average between invaded non invaded. For the legend of species check the main text. “na”: missing values due to the collapse of the box because of mold

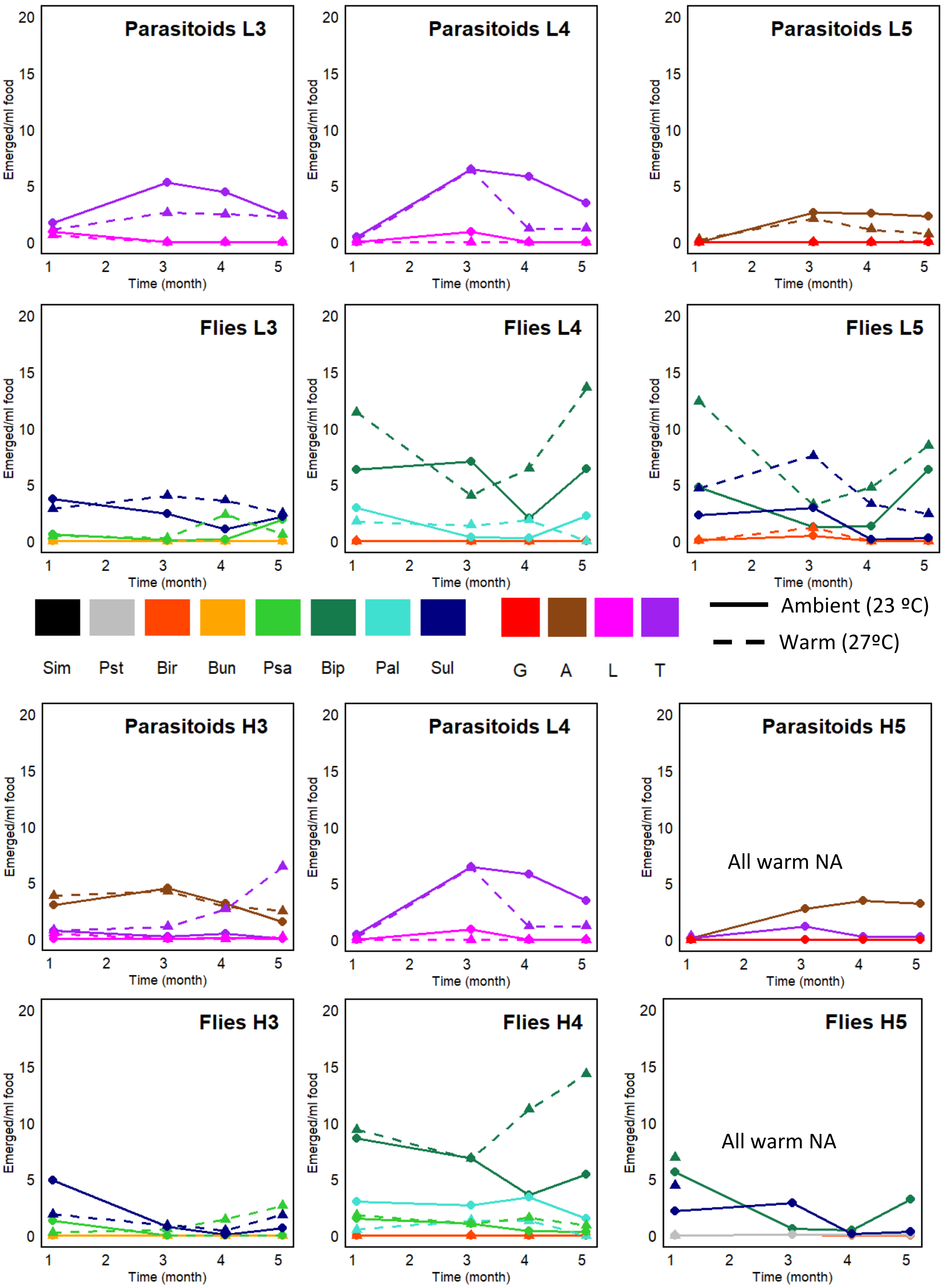


**Suplementary figure 1B.** Population dynamics (number of emerged adults/ml of food) of the different species in the food webs designs (L: low, H: high complexity) with no presence of *D. simulans* during 5 months. Solid line represent the boxes under ambient temperature (23 °C) and dashed line under warming (27°C). The data show the average between invaded non invaded. For the legend of species check the main text. “na”: missing values due to the collapse of the box because of mould

**Table S1** T-values of the quasi binomial model of Parasitism rate and quasipoisson model of abundances at different sampling times. The colours represent the level of significance (see legend below the table).

| | Emerged flies | | | | Emerged parasitoids | | | | Parasitic rate | | | |
|---|---|---|---|---|---|---|---|---|---|---|---|---|
| | **m1** | **m3** | **m4** | **m5** | **m1** | **m3** | **m4** | **m5** | **m1** | **m3** | **m4** | **m5** |
| **Warm** | 0.8 | 0.5 | 4.4 | 2.3 | 0.4 | -0.1 | -2.0 | -0.1 | 0.2 | -1.1 | -4.2 | -1.7 |
| **Initial complexity** | 0.9 | 0.2 | -0.8 | -0.1 | -0.9 | -0.3 | 0.8 | -0.7 | -0.8 | -0.2 | -0.7 | -1.0 |
| **Invasion** | -0.2 | 1.8 | 1.7 | 0.4 | 0.3 | 0.9 | 1.2 | 0.7 | 0.0 | -1.6 | -2.2 | -1.6 |
| ***D. simulans*** | 2.6 | -1.2 | -2.3 | -2.8 | 0.4 | 4.9 | 2.8 | 2.4 | -0.5 | 2.8 | 3.4 | 4.2 |
| *W27ºC:D.sim* | -1.8 | -1.6 | 0.2 | 2.3 | 1.2 | -1.3 | 2.3 | 0.5 | 1.1 | 1.4 | 0.3 | -2.2 |
| **p-value** | | < 0.05 | | 0.01 | | <0.001 | | | | | | |

**Table S2** Statistics (t-value) of the main predictors of the linear model of Inverse of covariance. The colours represent the level of significance (see legend below the table).

| | ICV | | | | | |
|---|---|---|---|---|---|---|
| | **Total** | **Flies** | **Parasitoids** | **Parasitism Rate** | **Shannon** | **Eveness** |
| **Warm** | 2.2 | 3.7 | -0.5 | 0.7 | -0.5 | 1.0 |
| **Initial complexity** | -0.5 | -0.1 | 0.5 | -0.6 | -1.0 | -0.1 |
| **Invasion** | 1.2 | 1.1 | 0.4 | -1.0 | 0.7 | 1.4 |
| ***D. simulans*** | -0.1 | -6.8 | -2.1 | 0.8 | -3.9 | -4.3 |
| **p-value** | < 0.05 | <0.01 | <0.001 | | | |
